# PathoClock and PhysioClock in mice recapitulate human multimorbidity and heterogeneous aging

**DOI:** 10.1101/2021.10.17.464755

**Authors:** Shabnam Salimi, Christina Pettan-Brewer, Warren C Ladiges

## Abstract

Multimorbidity is a public health concern and an essential component of aging and healthspan but understudied because investigative tools are lacking that can be translatable to capture similarities and differences of the aging process across species and variability between individuals and individual organs. To help address this need, body organ disease number (BODN) borrowed from human studies was applied to C57BL/6 (B6) and CB6F1 mouse strains at 8, 16, 24 and 32 months of age, as a measure of systems morbidity based on pathology lesions to develop a mouse PathoClock resembling clinically based Body Clock in humans, using Bayesian inference. A mouse PhysioClock was also developed based on measures of physiological domains including cardiovascular, neuromuscular, and cognitive function in the same two mouse strains so that alignment with BODN was predictable. The results revealed between- and within-age variabilities in PathoClock and PhysioClock, as well as between-strain variabilities. Both PathoClock and PhysioClock correlated with chronological age more strongly in CB6F1 than C57BL/6. Prediction models were then developed, designated as PathoAge and PhysioAge, using regression models of pathology and physiology measures on chronological age. PathoAge better predicted chronological age than PhysioAge as the predicted chronological and observed chronological age for PhysioAge were complex rather than linear. In conclusion, PathoClock and PhathoAge can be used to capture biological changes that predict BODN, a metric developed in human, and compare multimorbidity across species. These mouse clocks are potential translational tools that could be used in aging intervention studies.

## Introduction

An increase in the population of older adults comes with a rise in age-related health conditions [1]. With the increase in lifespan in the population, healthspan has become a focus of research and public health policies [2, 3]. Therefore, measurements of healthspan require cross-species translatable tools for preclinical and clinical studies [4–6]. For example, with distinct frailty indices in humans, mouse models of frailty have been developed [7–9]. Recent studies of age-related pathology using a geropathology platform that generates age-related lesion scores [10] have suggested a stronger correlation between age-related pathologies and chronological age than frailty indices [5, 10, 11]. A crucial aspect of healthspan is the burden of multimorbidity, conventionally described as having two or more diseases [2]. While comorbidity is commonly used to assess clinical disease burden in people, especially with increasing age, it has not been well characterized in animal models in a manner that has significant translational relevance.

Studies of rodent models have extensively focused on lifespan, using either chronological age or time to death as outcomes [12–14]. In recent years, healthspan has become an increasingly important focus of research with an increase in the older population [11, 15]. Multimorbidity is one of the crucial aspects of healthspan[4], but study of multimorbidity in mammalian models has been limited. Most studies have used a frailty index (FI) of one kind or another as a translatable tool to report health in humans or mice. Previously, FIs were adapted to apply to C57BL/6 mice [8], and a study of C57BL/6 showed that FI scores were related to heart hypertrophy[9]. However, the disease status of organs with morbidity and the histopathological changes associated with specific organs were not studied. In addition, whether the physiological changes tracked with end-point pathologies was not reported. Moreover, human FIs usually incorporate the disability state into the score, skewing the measures toward those with disability rather than predicting disability as one possible deteriorating outcome prior to mortality [7].

One approach to better define comorbidity in animal models is to consider the presence of pathology at the organ level. While many studies have focused on how aging and age-related diseases affect individual organs, each organ’s contribution to overall aging has been overlooked. A recent study of multimorbidity in the human population has suggested body organ disease number (BODN) as an index of multimorbidity [16]. The disease levels of each organ heterogeneously incorporate into BODN at the individual level. The integrated burden of disease incorporated into BODN for an individual has been shown to outperform chronological age to predict BODN and has been termed Body Clock [16]. Therefore, it is speculated that an organ-based pathology system in aging mice, such as the recently developed geropathology grading platform [10] could be used to define a measurable phenotype designated as PathoClock. By applying the Bayesian inference [9], the mouse-specific PathoClock could be a useful tool to simulate the human Body Clock. In addition, physiological and functional measurements are routinely determined in aging mouse studies. Therefore, this type of preclinical data could be used to predict heterogeneous BODN resulting in a mouse-specific PhysioClock.

Some studies of aging have used biological measures tied to chronological age as an outcome to predict biological age [12]. The current manuscript introduces PathoAge and PhysioAge using Bayesian inference and regression models of pathology and physiology measures, respectively, to understand how pathology and physiology based on chronological age align with biological age.

## Material and Methods

### Mice and study design

CB6F1 and C57BL/6 male mice were obtained from the National Institute on Aging (NIA) aged rodent facility (Charles River^@^ Laboratories) and housed in a specific pathogen-free facility at the University of Washington (UW) School of Medicine. Standard care procedures were followed including rodent chow, reverse osmosis purified automatic watering, 12:12 light cycle and 72±2 degrees F room temperature. All animal protocols were approved by the UW Institutional Animal Care and Use Committee. Animals were euthanized for pathology studies at age 8, 16, 24, and 32 months, three months after the physiological domains were measured.

### Physiological Assessment

#### Cardiac function

Echocardiography was used to assess systolic and diastolic function in mice. The Siemans Acuson CV-70 system using standard imaging planes: M-mode, conventional, and Tissue Doppler imaging, was used to measure cardiac function, including the ratio of the aorta and left atrium (AO/LA ratio), ejection time (ET msec), isovolumic contractile time (IVCT msec), isovolumic relaxation time (IVRT msec). The E/A ratio as a marker of the left ventricle function indicates the peak velocity blood flow from left ventricular relaxation in early diastole (the E wave) to peak velocity flow in late diastole caused by atrial contraction (the A wave). Myocardial performance index (MPI) incorporates both systolic and diastolic time intervals in expressing a global systolic and diastolic ventricular function quantified as MPI=(IVCT+IVRT)/ET [17]. The methods are described elsewhere [18].

#### Neuromuscular Function

Established tests of muscular activity were used to assess changes in muscle strength and coordination with age. Several assessments including coordinated walking ability, grip strength, novel environment response, and self-motivated running, were used to address variability due to motivation, emotionality, or sensory deficits.

##### a. Coordinated walking ability

Coordinated walking ability was assessed using a rotarod apparatus (Rotamax 4/8, Columbus Instruments, Inc.) that tested the ability of the mouse to maintain walking speeds on a rotating rod. Mice were placed in the lanes of the rotarod with initial rod speed set at 0 RPM. The speed was progressively increased by 0.1 RPM/sec (0 to 40 RPM over 5 minutes) until all mice had been dislodged as determined by an infrared sensor. The time in seconds was recorded. Three successive runs were performed per day for three days. Therefore, there is an evaluation of motor function and performance learning. The assay was performed by the same person, at the same location. Data are reported as the median of 3 trials and standardized by body weight.

##### b. Grip strength

Forelimb grip strength was analyzed using a force tension apparatus (San Diego Instruments Columbus Instruments, Inc.). Prior to the test, each mouse was weighed to the nearest 0.1 g. Once mice gripped the stationary bar with their forepaws, they were stretched horizontally while held at the base of their tails. Mice were pulled gradually until they let go of the bar. The process was repeated 5 times to determine the peak grip force value (gram-force) standardized to body weight[19].

##### c. Novel environment response

Mice were assessed for movement levels in a novel cage environment using an open field photobeam apparatus (Photobeam Activity System, Columbus Instruments, Inc.). Each mouse was placed for five minutes in a clear, rectangular, plastic container the size of a standard mouse cage, which had a rectangular grid of infra-red beams inside, three on the X axis and four on the Y axis to measure horizontal movement (lateral activity). Another grid set of beams was positioned above the lower set to measure vertical movement (rearing). Beam breaks were counted for activity and rearing, and further classified for either the central or peripheral part of the box as a measure of anxiety. Activity was assessed for a five-minute period on three consecutive days[20].

##### d. Self-motivated running

Self-motivated running distance was measured by a voluntary wheel running apparatus over three days as described by Goh and Ladiges (2015)[21]. Mice were placed into a standard cage with a slanted plastic saucer-shaped wheel (Med Associates, Inc.). Mice were acclimated to the cage for 48 hours with the wheel locked, after which the wheels were unlocked and the distance each mouse ran was tracked by a computer over 72 hours including both light and dark cycles. Total distance in kilometers was recorded.

#### Cognitive Function

Cognition was assessed using the radial water tread (RWT) maze, an assay used to assess memory as previously described [22, 23]. The RWT detects changes in hippocampal function in mice. Briefly, mice are introduced into an approximately 30- inch circular galvanized enclosure with waste-deep water and peripheral escape holes in the sides at regular intervals, all closed except one which leads to a dark “safe box” with a heating pad. The inside walls contain spatial cues for the animal to find the escape route with repeated trials. The animals were given three trials per training day, and the testing period ran across successive days to test long-term memory acquisition. Performance was recorded by direct observation [24].

Various physiological domains described above were used to predict body organ disease number (BODN) and define a mouse-specific PhysioClock independent of chronological age.

### Pathological Assessments

#### Cataract assessment

The presence and severity of cataracts were assessed by slit-lamp ophthalmoscopy on unanesthetized mice after dilation with a 3:1 volume mixture, respectively, of tropicanamide and phenyl hydrochloride to achieve full dilation. The degree of lens opacity was rated by half steps from 0 (completely clear) to 4 (complete opacity of a mature cataract) as previously described[25].

#### Histopathology assessment

Histopathology assessments were performed on Hematoxylin and Eosin-stained 4-micron tissue sections from heart, kidney, liver, pancreas, muscle, lung, and brain as previously described [10]. Age-related lesion severity levels were determined by a scoring system from 0 (no lesion present) to a range of 1 to 4 (lesion presence and severity). The absence or presence and severity of age-related lesions were then used to determine organ morbidity defined as the presence of two level 1 lesions or one lesion with a score of 2 or greater. The body organ disease number (BODN) was then calculated as the number of organ systems with morbidity as a proxy of multimorbidity and a counterpart of clinically determined BODN in humans. With the premise that different pathology entity levels incorporate into BODN heterogeneously, all organ pathologies in a model were used to predict BODN for each mouse to quantify PathoClock independent of chronological age.

## Statistical Analyses

BODN was considered an ordinal outcome as a number of organ systems with at least two positive pathology criteria at level 1 or at least one pathology at level 2 or more. We recorded the levels starting from 1 as an ordinal value. Bayesian inference was used for ordinal outcome [26] also ordinal[27], binary, or continuous predictors depending on the type of predictor variables [27–30]. Bayesian inference approach was used comprising two components: 1) Prior knowledge on the estimates (parameters), as the information before observing the data P(⍰) where ⍰ indicates the parameters; and 2) the likelihood P(Y|⍰) of the information contained in the data (Y). Using the Bayes formula, the posterior distribution of the parameters P(⍰|Y) was obtained, which can be updated when encountering new data[28].

Applying the conditional probability given the known data on BODN, the Bayesian inference framework yields the posterior density of beta estimate coefficients and 95% credible interval (CI) that each pathology level incorporates into BODN, or each physiological measure predicts BODN. For each coefficient parameter we determined the distributions of their prior parameters using weak priors[28]. For the classes of beta coefficients and intercept, the prior estimate with a normal distribution (mean: μ = 0, variance σ = 10), and for the class standard deviation (sd) which indicated the variation of levels related to varying age, the half-Cauchy (0,10) was used. The uniform prior with a Dirichlet distribution was used for ordinal predictors (i.e., (2, 2) for ζ1 and ζ2 [simplex parameters] for a three-level pathology predictor). We reported the standard deviation for the model level in multilevel analyses (sd), and sigma which is the variance of a continuous outcome in the model with gaussian family[27, 30]. *Posterior predict* function was then used in the Bayesian framework [28, 30] to predict individual-based BODN for each mouse using all organ pathology levels, termed PathoClock. The correlation of PathoClock and chronological age was quantified as a rate of pathology-based biological age.

The model accuracy was assessed with leave-one-out cross-validation (LOO-CV) (k<0.7) which with a Pareto-smoothed importance sampling diagnostic k <0.7 indicating the LOO-CV computation is reliable and there are no outlier observations. Also, a Bayesian inference approach called “stacking” determines model weights for each model to predict an outcome [29, 31]. The Leave-One-Out R squared (LOO_R^2^) was used to determine the R^2^ of the model to show how a model explains the outcome[32]

All statistical analyses were performed using the Bayesian “brms” software package[30]. A dynamic Hamiltonian Markov chain Monte Carlo (MCMC) algorithm [30, 33] was used to obtain posterior draws using a minimum of six chains and a minimum of 10,000 iterations. Model averaging was then used in the Bayesian framework, called stacking, which provides weight for the best model predicting BODN[34]. For pathologies more than two levels we used “monotonic” effect implemented in the Bayesian inference framework which defines probability of coefficient estimates with Dirichlet distribution[27]. The physiological measures like grip strength or rotarod, which have an inverse association with BODN, were transformed so that were multiplied by −1 to develop PhysioClock.

Commonly, studies statistically have regressed biomarkers or phenotype measurements on chronological age to assess how they predict chronological age [12]. In concert with such an approach, we developed PhysioAge and PathoAge, regressing the allocated physiological measures and pathological level measurements, respectively, on chronological age using gamma distribution. In CB6F1, to develop PhysioAge, we included normalized grip strength, rotarod test at day 3, open field activity at day3, open field rearing at day 2, distance, AO/LA ratio, ET (ejection time), LVM, MPI, and Maze test at day 5. In C57BL/6, for PhysioAge we included nine physiologic measures including AO, LA, natural log transformed E/A ratio, LVMI and MPI, Maze test at day 5, open filed activity at day 1, rotarod at day2, and normalized grip strength.

## Results

### Physiological performance predicts body organ disease number

#### Cardiac function

Echocardiography was used to measure cardiac function. For CB6F1, the ratio of aortic valve diameter to left atrium dimension (AO/LA) was inversely associated with increase in BODN (beta=−2.3, 95% CI: −4.3 to −0.32), E to A waves ratio (beta=−1.5, 95% CI: −2.9 to −0.15), on natural logarithm scales and with relatively high uncertainty (wide credible interval CI) isovolumic contraction time (IVCT) [beta=1.48, 95% CI: 0.02-3.0], left ventricular internal diameter end diastolic (LVIDd) [beta=7.6, 95% CI: 0.13=15.0], left ventricular internal diameter end systolic (LVIDs) (beta=4.7, 95% CI: 0.47-9.03), (considering both systolic and diastolic measures (MPI) [beta=2.03, 95% CI=0.11- 4.01], ejection time (beta = − 6.6, 95% CI: −12.0 to −0.7) predicted BODN (Figure 1A). Chronological age per se was strongly associated with BODN (beta=0.34, 95% CI=0.24-0.46) while stacking of the cardiovascular models predicting BODN revealed cardiovascular physiologies predict BODN stronger than chronological age predicted BODN so that the model weight for chronological age turned to zero. The largest model weights were allocated to ejection time (ET: 27.5%), E/A ratio (26%), IVCT (21.5%), MPI (17%), AO/LA ratio (8%) that MPI by 46%, LVIDd by 19%, E/A ratio by 28.5%, IVCT (6%) with the rest also turned to zero.

**Figure 1.**
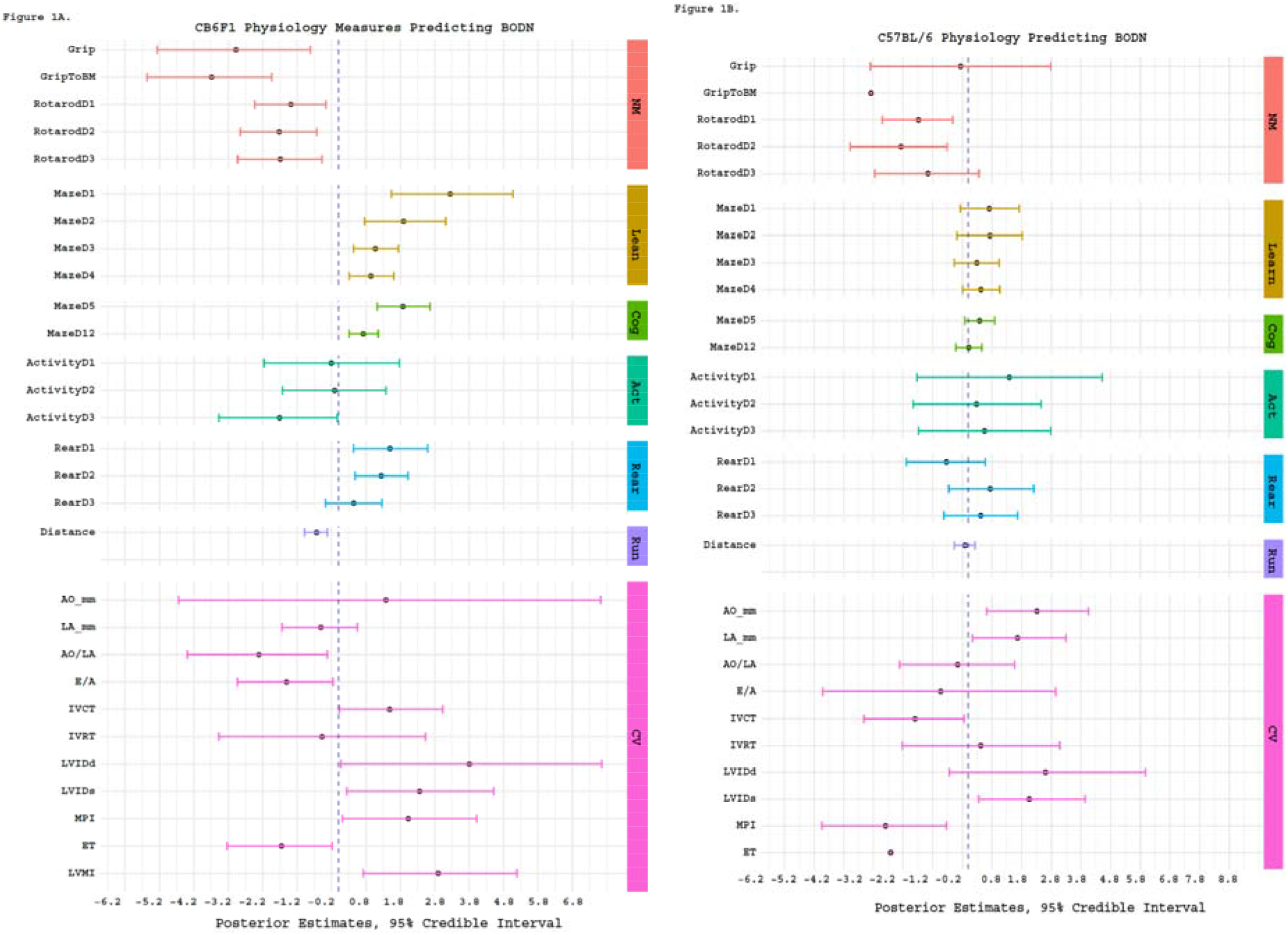
Physiologic predictors of Body Organ Disease Number (BODN) in **A)** CB6F1, **B)** C57BL/6 mice. Grip: Grip strength, GriptoBM: normalized grip strength to body size, RotarodD1: rotarod at day 1, MazeD1: Barnes maze at day1, ActivityD1: Open Filed activity at day1, Rear: Open filed activity rearing, AO: aortic valve dimension in millimeter, LA: left ventricular valve, AO/LA: the ratio of AO to LA dimensions, E/A, E wave to A wave ratio, IVCT: isovolumic contractile time millisecond (msec), IVRT: isovolumic relaxation time (msec), LVIDd: left ventricle internal diameter end diastole, IVIDs: left ventricle intra diameter end systole, MPI: myocardial performance, ET: ejection time., NM: neuromuscular, learn: learning stage, Cog: cognition, Act: Open filed activity, Rear: Open filed activity rearing, Run: Voluntary wheel running, CV: Cardiovascular physiology.

For C57BL/6 mice, aortic valve diameter (beta=2.3, 95% CI: 0.6-4.0), LA (beta=1.65, 95% CI: 0.13-3.28), IVCT (beta=−1.79, 95% CI: −0.35 to −0.14), LVIDs (beta=4.1, 95% CI: 0.69-7.8) were significantly predicted BODN. With wide uncertainty (credible interval including zero) both AO/LA ratio and E/A ratio were inversely predicted BODN.

Using model averaging over all physiological measures of the cardiovascular system in C57BL/6 showed that MPI by 72.6%, LVIDs by 22%, E/A ratio by 0.5%, aortic diameter (mm) by 1 %, and left atrial diameter (mm) by 4.5 % predicted BODN and were included in the final model as cardiovascular physiology domains to quantify PhysioClock.

#### Neuromuscular Function

Rotarod test for CB6F1 indicating disturbed balance state predicted increase in BODN measured at day 2 (beta=−1.7, 95% CI: −2.86 to −0.62). Stacking of the models showed that the model assessed balance state at day 2 weighed 75.4% to predict BODN compared to day 1 (17.1%) and day3 (7.5%). Therefore, we included rotarod test day 2 in the PhysioClock model. The lower the grip strength, the larger the BODN was (beta=−5.9, 95% CI: −10.5 to −1.6), and it was more robust when normalized to body mass (beta=−7.3, 95% CI: −11.4 to −3.8). Comparing models showed that the normalized grip strength over body size was a stronger predictor of BODN at 98.3%.

For C57BL/6 mice, balance states at day 1 and 2 significantly predicted BODN with day-2 model weight (53.2%) larger than day 1 (46.8%). Therefore day 2 was included in the Physiology Clock. Like CB6F1, the grip strength in C57BL/6 normalized over body mass was a stronger predictor of BODN (Figure 1B). In this strain, only grip strength normalized by body mass significantly predicted BODN with credible intervals excluding zero. However, wide CI (beta=−3.2, 95% CI=−6.4 to −0.18) showed some degree of uncertainty.

#### Cognitive Function

For CB6F1 mice, time of learning maze measured at day 1, 2 ,3 and 4 was associated with increased in PathoClock (day1: beta=1.13, 95% CI=0.18-2.07; day2: beta=0.87, 95% CI=0.21-1.54; day4: beta=0.45, 95%CI=0.11-0.78). The longer the learning process at day 1 the larger the PathoClock was. The longer maze test indicated poorer cognition and predicted larger PathoClock (day 5: beta=0.49, 95% CI=0.15-0.83; day 12: beta=0.92, 95%CI:0.49-1.38). Likewise, results were detected for observed BODN (day2: beta=1.5, 95% CI: 0.22-2.79; day4: beta=0.62, 95%CI: 00.08-1.18) and this association was stronger at day 2. Also, cognitive decline was associated with increase in BODN so that the maze test results at day 5 (beta=0.94, 95% CI= 0.35-1.53) and 12 were strongly predictive of BODN (beta=0.92, 95%CI:0.49-1.38). The model stacking over the models including learning stages showed learning stage at day 1 (weight 62.8%) was a stronger predictor of BODN, and the cognition test at day 5 was stronger than day 12 (weight by 98.6%). Stacking over the learning and cognition test stage models showed that day 1 and day 5 weighed more than other days (model weights for day 1: 19.6%, and day 5: 78.9%). We included these two measures of learning and cognition in the PhysioClock model for CB6F1.

For C57BL/6 mice among learning and cognitive stages of the RWT maze test, day 1 and day 5 were the more robust predictors of BODN with model weights 61.5% and 99%, respectively. However, overall, the maze test in C57BL/6 was less predictive of BODN compared to CB6F1, but we included the day 5 maze test as it carried a larger weight to predict BODN.

#### Open field physical activity, rearing, and wheel running distance

For CB6F1 mice, open field physical activity indicating physical aptitude at day 3 significantly and inversely predicted BODN (beta=−1.7, 95% CI: −3.4 to −0.03). The higher rearing in physical activity, the larger the BODN was with a larger estimate at day 1 (beta=1.49, 95 CI: 0.44-2.59). However, the model weight favored the rearing activity at day 2 in prediction BODN weighed by 77.2% compared to day 1 (22.8%). We included day 2 rearing activity in the final model determining PhysioClock. The mice with lower running distance had larger system morbidity measured by BODN (beta=−0.6, 95% CI: −0.98 to −0.3).

For C57BL/6 mice, open field physical activity was not significantly predictive of BODN. Stacking the models showed day 1 open field activity model weight was 79.8%. Also, the rearing activity model at day 1 with 57.4% weight explained BODN better than day 2 and 3. The total distance for the running wheel was inversely associated with BODN yet with a broad uncertainty (beta=−0.11, 95% CI=−0.46 to 0.23). We only included total distance in the final model to quantify PhysioClock.

### Organ Pathology heterogeneously integrates into body organ disease number

Overall, the BODN was higher for CB6F1 than C57BL/6 at any age. The median and range of BODN were 2 (1-4) at 8 months, 4 (2-5) at 16 months, 5 (4-7) at 24 months, 6 (5-7) at 32 months for CB6F1; and for C57BL/6 they were 3 (2-4) at 8 months, 5 (4-6) at 16 months, 5 (4-7) at 24 months, 6(4-7) at 32 months.

Because the pathologies are ordinally graded based on severity, we used the monotonic effect that is applicable when the levels are not equidistant, and we showed that each organ’s pathology severity scores heterogeneously incorporated into body organ disease number (BODN). The result shows that the degree of organ pathology heterogeneously incorporates into BODN in both B6F1and C57BL/6 mice and some pathologies are more dominant for each strain (Figure 2A-B).

**Figure 2.**
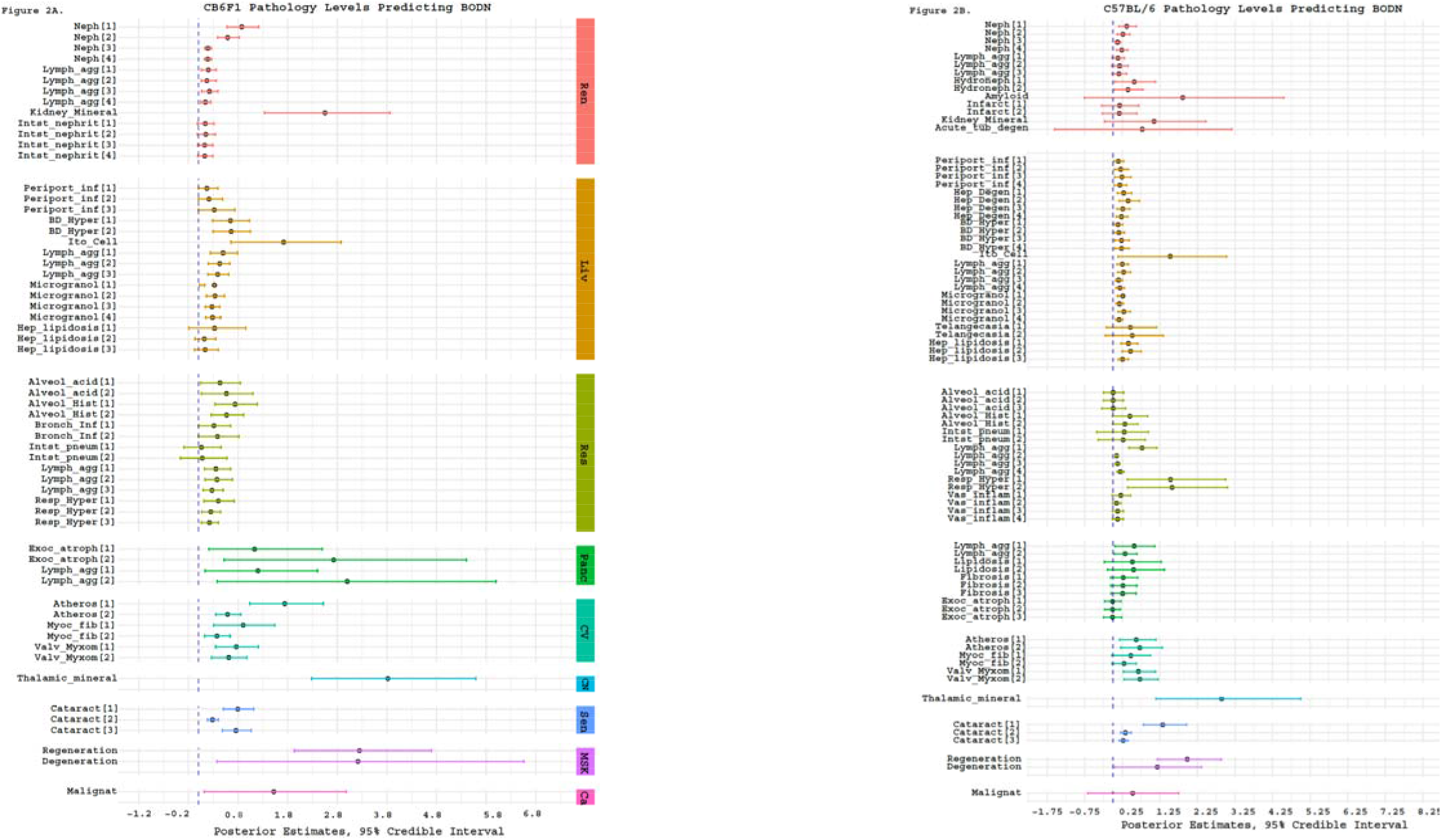
Pathology levels of seven organs in **A)** CB6F1 and **B)** C57BL/6 male mice incorporating Body Organ Disease Number. Abbreviations: Neph: nephropathy, lymph_agg: lymphocyte aggregation, Infarct: infarction, Kidney_mineral: mineral deposition in kidney, Acute, tub_degen: acute tubular degeneration, perioport_inf: periportal infiltration, Hep_Degen: hepatic degeneration, BD_Hyper: bile duct hyperplasia, Ito_cell: itocell hyperplasia, Microgranol: microgranuloma, Hep_lipidosis: hepatic lipidosis, Alveol acid: alveolar macrophage pneumonia, alveol_hist: alveolar histiocytosis, Resp_hyper: respiratory duct hyperplasia, Vas_inflam: perivascular inflammation, exoc_atroph: exocrine atrophy, Atheros: atherosclerosis, Myco_fib: myocardial fibrosis, Valve_Myxom: valvular myxomatosis, thalamic mineral: mineral deposition in thalamus area, Regeneration: skeletal muscle degeneration, regeneration: skeletal muscle regeneration, malignant: malignant cancer.

We included all pathologies that predict BODN with high accuracy using LOO-CV (k<7). The estimates of each pathology level incorporating into BODN are depicted in Figure 2A and B. The complete model including all organs’ pathology to predict BODN explained variability of BODN by 87% and 88% for C57BL/6 and CB6F1, respectively. We quantified PathoClock from the model, including all organ systems to predicted BODN using age as levels. There is inter-mouse variability of PathoClock even within the same chronological age. Mean PathoClock was mainly larger in CB6F1 compared to C57BL/6, especially at age 28 months (6.5±1.10 vs. 6.3±0.64) and 32 months (7.6±1.5 vs. 7.3±0.65), respectively. The between-strain variability (variance) over the age spectrum was 3.3 months. In CB6F1, cardiovascular-related pathologies with higher uncertainty (the narrow credible intervals excluding 0) were significantly incorporated into BODN (Figure 2A).

While specific pathologies of each organ variably incorporate into BODN of the renal system, only kidney mineral disposition had a wide uncertainty in CB6F1. In the C57BL/6 mice, in addition to mineral disposition, amyloid accumulation and acute tubular damage had wide credible intervals and large uncertainty predicting BODN (Figure 2B). In C57BL/6, the majority of liver-related pathologies were incorporated into BODN yet heterogeneously. Of note, incorporation of the regeneration state in BODN in C57BL/6 mice was larger than of the degeneration state. Interestingly, lymphoid aggregates in almost all organs significantly incorporated into BODN.

### Correlation of PathoClock, PhysioClock, and chronological age is strain dependent

To understand how well the two final models, the one including all pathology levels to develop PathoClock and the one including physiology measures to quantify PhysioClock, explained BODN, we used the Leave-One-Out R-squared (LOO_R^2^) method. The model including all pathologies to predict BODN for C57BL/6 (PathoClock), explained about 87% of BODN (LOO_ R^2^=0.87), while the model used to develop PhysioClock explained BODN by 64% (LOO_R^2^=0.64). For CB6F1 mice the models to develop PathoClock explained BODN by 94% (LOO_R^2^=0.94), and the model used to develop PhysioClock explained BODN by 67% (LOO_R^2^=0.67). In both strains the models from which PathoClock was extracted explained BODN better than PhysioClock; however, in CB6F1 the overall model performance was better than in C57BL/6.

The distributions of PathoClock and PhysioClock are depicted in interactive Figure 3. The correlation of PathoClock and chronological age was r=0.75 in C57BL/6, but in CB6F1 the correlation was larger (r=0.80) (Table 1). In some individual mice, PathoClock was smaller than BODN, while in some mice, it was larger. These results suggest variability in incorporating pathology levels in the same age group and across ages. Also, a larger impact for pathology levels can be manifested as a larger PathoClock in mice within the same age group or in an older group. In contrast, a smaller PathoClock at an older age may suggest minor impact of pathology levels on BODN despite an older age. This result opens a roadmap to study resilience and body system reactions in relation to pathology (Figure 3A-B). Correlation between PhysioClock and age at euthanasia in CB6F1 mice was r=0.71 with variability across age groups by 3.72 months (sd=3.7), and the correlation of PhysioClock with age at euthanasia was r=0.68 for C57BL/6 with 6.5 months variability in age (sd=6.5; Table 1). Some mice with higher BODN had lower PhysioClock (at middle age), suggesting physiological resilience to the development of pathology. However, it could also be due to an insufficient adaptation response. Despite the high correlation, the patterns of both PathoClock and PhysioClock in relation to chrononlogical age were not linear, and exponential patterns were detected (Figure 3 C-D).

**Figure 3.**
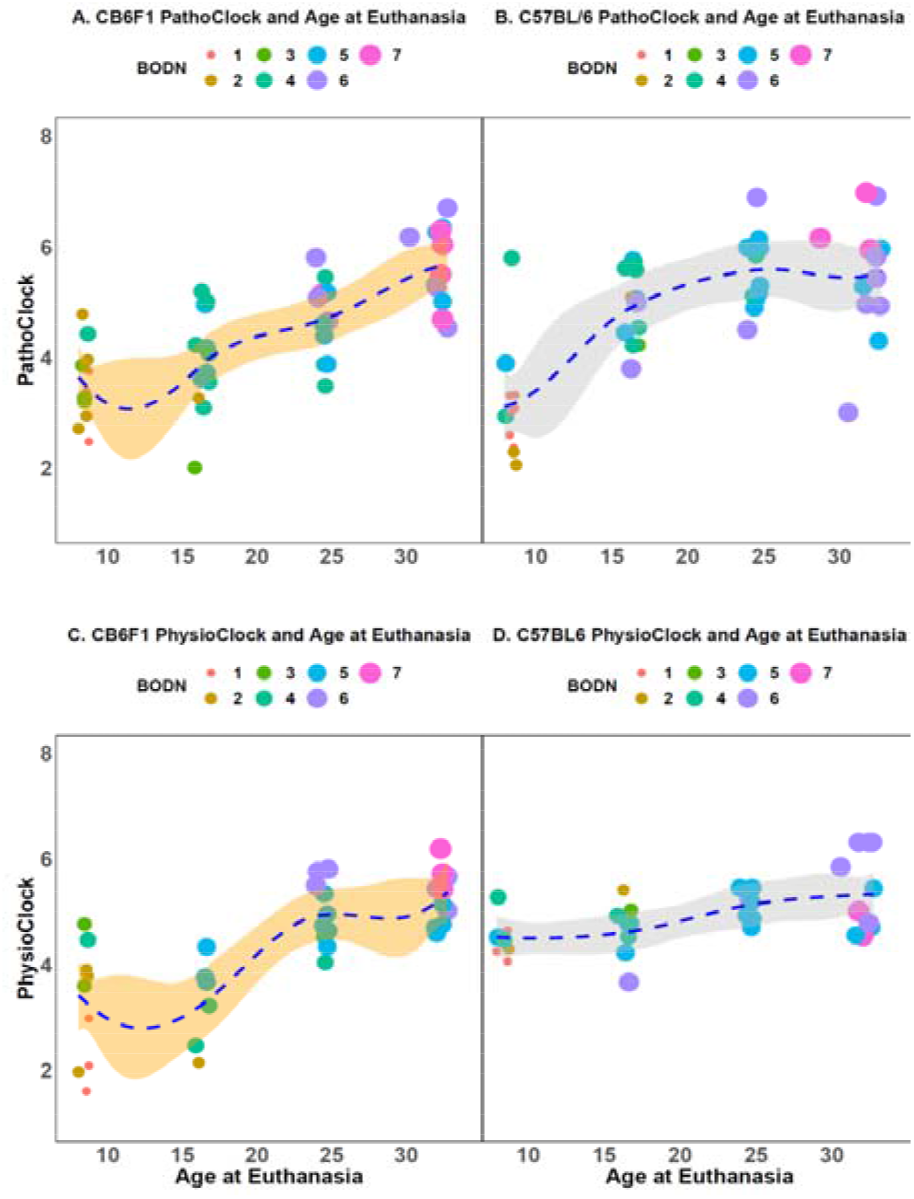
The distribution of PathoClock and PhysioClock by age. To see the 3D figures, click on the included links. PathoClock is determined by how each pathology level incorporates into Body Organ Disease Number (BODN). **A. PathoClock in CB6F1 mice and age at euthanasia. B. PathoClock in C57BL/6 mice and age at euthanasia.** PhysioClock was determined by how each physiological measures predicted BODN. **C. CB6F1 PhysioClock and age at euthanasia. D. C57BL/6 PhysioClock and age at euthanasia.**

**Table 1.** Leave-One-Out (LOO) R squared of models for developing PathoClock, PhysioClock, PathoAge and PhysioAge and the correlation with chronological age. Sd: standard deviation for model levels in multilevel analyses, sigma is the variance of gaussian family for continuous outcome.

|  |  |  |  |  |
| --- | --- | --- | --- | --- |
|  | LOO-R<sup>2</sup>, | Correlation (r) | LOO-R<sup>2</sup> | Correlation (r) |

|  | (Sd or sigma)<br>C57BL/6 | with ChAge<br>(C57BL/6) | (Sd or sigma)<br>CB6F1 | with ChAge<br>(CB6F1) |
| --- | --- | --- | --- | --- |
| Model for PathoClock | 0.87,<br>(sd=5.7 months) | 0.76 | 0.94,<br>(sd=7.4 months) | 0.80 |
| Model For PhysioClock | 0.64,<br>(sd=6.7 months) | 0.68 | 0.67,<br>(sd=3.7 months) | 0.73 |
| Model for PathoAge | 0.86,<br>(sigma=2.5 months) | 0.98 | 0.93,<br>(sigma=2.4 months) | 0.98 |
| Model for PhysioAge | 0.75,<br>(Sigma=3.9 months) | 0.83 | 0.70,<br>(Sigma=4.0 months) | 0.93 |

### PhyisoAge and PathoAge align with chronological age in a strain dependent manner

The common approach to measuring the rate of aging with chronological age has been to regress phenotype measurements over chronological age. Likewise, we developed PhysioAge and PathoAge by regressing the physiological and pathological measurements on chronological age and determining the R^2^ of the model using the Leave-One-Out approach (LOO_R^2^) and then assessing the correlation between predicted age and chronological age for PathoAge and PhysioAge. In C57BL/6 mice, PathoAge, having a larger LOO_R^2^ and stronger correlation with chronological age, explained chronological age better than PhsyioAge. PathoAge variability across age was 2.5 months while variability of PhysioAge was 3.9 months (Table 1). In CB6F1, both PathoAge and PhysioAge strongly explained chronological age, with PathoAge (LOO_R^2^=0.93) explaining chronological age better than PhysioAge (Loo_R^2^=0.7). In C57BL/6 mice, there was a slow slope of correlation between PhysioAge and ChAge so that the PhysioAge at younger ages had similarity with middle age groups (Figure 4).

**Figure 4.**
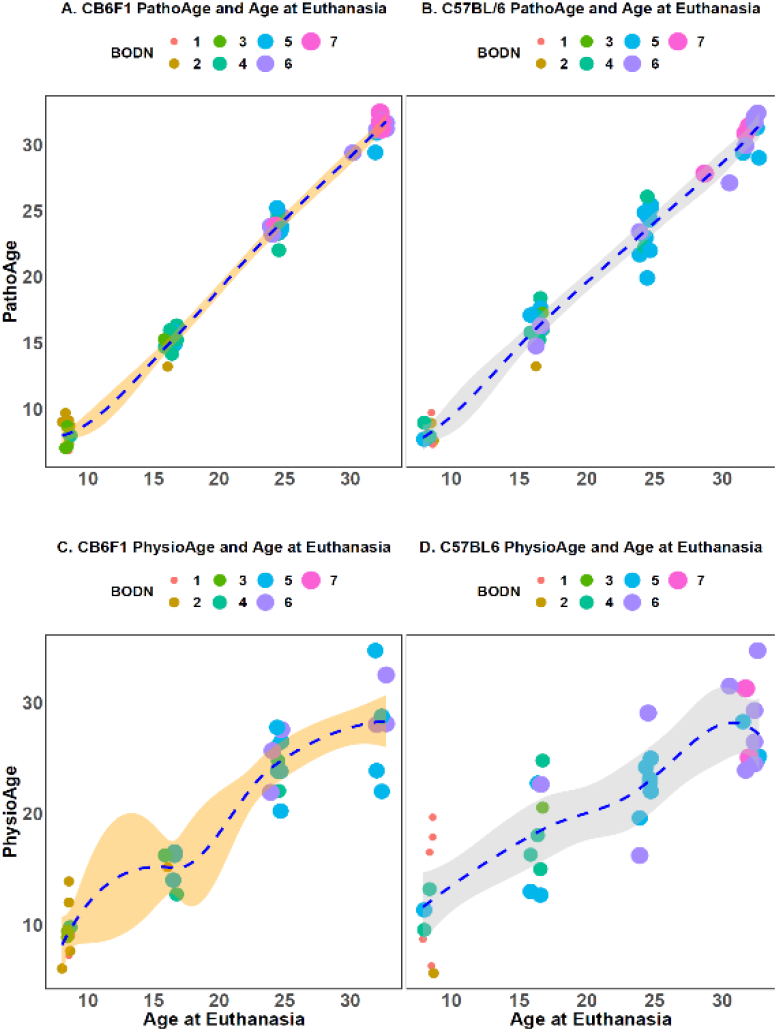
Developing. developed PathoAge pathologies were included in a model to predict ChAge in both **A.** CB6F1 and **B.** C57BL/6. Developing PhysioAgek, the same physiological measures as the ones used in Physioclock, were regressed over ChAge in both **C.** CB6F1 and **D.** C57BL/6. While PathoAge are almost linearly predict ChAge with some subtle degree of uncertainty, PhysioAge in both strains endures more uncertainty to predict ChAge. The size and color indicate the increase in number of body organ disease number (BODN)

## Discussion

In this study, physiology performance and pathology data were generated from C57BL/6 and CB6F1 male mice ranging from 4 to 28 months of age. As a result of these data, pathology-based multimorbidity as an outcome was developed and is reported for the first time, with the pathological and physiological determinants designated as PathoClock, and PhysioClock, respectively. Using histopathology lesion scores in each organ as a proxy for diseases, the morbidity of each organ system was defined as at least two low pathology grades (=1) or one higher pathology grade (>1). The sum of the organ systems’ morbidities determined the Body Organ Disease Number (BODN) as a new outcome representing a global index of health at body organ system level, resembling what was recently developed and validated in a multimorbidity study of human aging [16]. The degree to which each organ-specific pathology level incorporates into BODN was assessed. The mouse strain-specific predicting BODN using pathology levels was termed PathoClock, a counterpart of Body Clock in humans [16]. Because physiological responses can vary by age and disease level, BODN was used as an outcome for determining physiological predictors developing PhysioClock which association with chronological age was assessed. The results showed that various levels of pathology of various organs heterogeneously incorporate into BODN. CB6F1 mice had a larger BODN and PathoClock compared to C57BL/6 mice in the same age group.

Interestingly, the two strains had distinct pathological and physiological components that predicted BODN. While aortic valve (AO) and left atrium (LA) dimensions significantly predicted BODN in C57BL/6 mice, in CB6F1 mice only the AO to LA ratio was a significant predictor of BODN. There was an inverse association of the E/A ratio with BODN in CB6F1. A decreased E/A ratio which is usually an indicator of diastolic heart failure suggests fibrosis so that the left ventricle cannot be filled with blood during the diastolic period between two contractions. Similarly, heart failure in humans is one of the age-related changes incorporated into BODN[16] and a health burden underlying hospitalization of older adults[35]. Moreover, in older adults decrease in E/A ratio incorporates into low exercise intolerance. The results in CB6F1 mice showed both an inverse association of voluntary exercise (running distance) and E/A ratio with BODN.

In both strains, while the Left ventricle dimension in end-systole (LVIDs) significantly predicted BODN, the left ventricle dimension in end-diastole (LVIDd) predicted BODN but with larger uncertainty. Shortening ejection time (ET), which has been suggested as a single indicator of human heart failure[36–38], significantly predicted BODN in CB6F1. Human study of echocardiographic measures has shown that a combination of both systolic and diastolic impairments is a better predictor of heart failure [37], as such a measure like the myocardial performance index (MPI) was a significant predictor of BODN in CB6F1, and it also predicted BODN in C57BL/6, albeit with some uncertainty. Left ventricular hypertrophy index normalized by tibial length (LVMI), an age-related change, significantly predicted BODN in both strains. Cardiac physiology markers were associated with BODN more strongly in CB6F1 mice than C57BL/6. Having more uncertainties in cardiac physiology measures, C57BL/6 mice might manifest cardiac physiology changes late in life or have physiological adaptation to histopathological changes later. Although PhysioClocks for both strains were associated with chronological age at euthanasia, the correlation was stronger in CB6F1, and there was larger variability in PhysioClock in C57BL/6 than in CB6F1. Replicative studies and response to interventions are required to replicate cardiac physiology changes in response to pathology.

There was variability in both organ physiology and pathology across strains and age groups. The ability to maintain neuromuscular and cognitive performance is an important component of healthspan in aging. Impaired physical activity and function are both cause and consequence of disease in human[39]. Albeit heterogeneous, older C57BL/6 mice had uncertainties in physical activity capacity in relation to BODN, while CB6F1 showed decreased balance, physical activity, lower running distance, and lower grip strength, all of which predicted increased BODN. All of these measures had a larger uncertainty in C57BL/6 to predict BODN. One possible explanation for wider uncertainty of physiologic measures in prediction of BODN in C57BL/6 is that some male C57Bl/6 mice at age 4 months might have already commenced physiological changes in response to pathology so that they are already similar to middle-aged mice. However, the C57BL/6 PhysioClock at older ages showed a relatively slow slope over the age spectrum which suggests resilience in physical function due to regenerative capacity in skeletal muscle in this strain as shown in their histopathology and association with BODN.

CB6F1 mice showed a more significant cognitive decline, attenuated volitional physical activity, disturbed balance, and diminished motor function in predicting BODN, while such functional measures did not significantly predict BODN in C57BL/6. The results suggest that C57BL/6 are also more resilient to functional decline than CB6F1 and/or might develop functional decline variability. While these two strains are commonly used in the study of normal aging, our results suggest strain-specific variability in pathological and physiological domains. However, mechanisms of functional resilience and whether there is more variability in functional impairment in C57BL/6, despite developing pathologies, can be explored comparing PathoClock and PhysioClock in both strains and measuring in-depth mechanistic markers in response to anti-aging interventions.

Recent reports in both CB6F1 and C57BL/6 mice show different organ aging, suggesting higher pathology scores in the cardiovascular system in CB6F1 and early onset of liver and kidney aging in C57BL/6 and organ-specific response to anti-aging interventions [40]. Because the aging kidney and liver show early and dominant age-related characteristics in C57BL/6, the inclusion of physiological markers of such organs to predict BODN may improve PhysioClock for both strains. Moreover, adding more organs to pathological studies, and obtaining more information by applying artificial intelligence to the images to extract high throughput information on echocardiography, pathology and other imaging can be incorporated into BODN and can update PathoClock and PhysioClock whenever this information is available.

In both strains, PathoClock was more strongly correlated with chronological age, with the CB6F1 PathoClock having a larger correlation, and we found variability in components of pathology and physiology across age groups. Recently, a new study applying the frailty component on chronological age FRAIL (Frailty Inferred Geriatric Health Timeline) and measuring lifespan with AFRAID (Analysis of Frailty and Death) in C57BL/6 mice predicted age with r^2^=0.64 in the test data [12]. Our models based on pathology or physiology more significantly predicted the animals chronological age with PathoAge in both strains and PhysioAge mainly in CB6F1. In addition, PathoClock based on BODN showed a stronger coorelation with chronological age (in CB6F1, r=0.8; in C57BL/6, r=0.76), and likewise, PhysioClock had larger correlation with chronological age (CB6F1: r=0.73; C57Bl/6: r=0.68). In both strains, the models from which PathoClock was extracted explained BODN better than PhysioClock, with model performances better for CB6F1 than C57BL/6. Similarly, PathoAge better predicted chronological age than PhysioAge with a larger correlation between observed and predicted chronological age. One possible explanation for the different correlations is the wider variability and uncertainty in physiological measures in predicting BODN and chronological age. Also, the data showed an exponential associations between physiological-based predicted age and chronological rather than linear. Another possibility is that we had smaller sample sizes for physiological measurements (30 mice in C57BL/6 and 35 in CB6F1). To better delineate pattern recognition, replication of these analyses including larger sample size would be helpful.

The two clocks developed, PathoClock and PhysioClock, are strong healthspan tools. In human aging, metrics that are statistically trained on phenotypes also predict health state [16]. One caveat of basing the data on chronological age is that there is arbitrarily consideration of chronological age as a variable outcome, while chronological age is a fixed number in an equation. Moreover, biomarker-based measures can fluctuate irregularly across age spectrums due to a variety of reasons such as adaptation, resilience, or severe organ damage. However, prediction of health outcomes like BODN can capture biological and pathophysiological changes independent of chronological age, as well as variability of biological age. While BODN and PathoClock can be used at the end point for healthspan, the PhysioClock can be used as a repeated measure in longitudinal studies to predict healthspan over time. The results of previously measured pathologies can be applied in the Bayesian models we developed, along with physiological measures, to predict BODN in aging studies using mice and can be used dynamically to further delineate mechanisms of aging [41]. Including components of pathology and or physiology into the models provides the ability to predict chronological age and integration into global health status measured as BODN. Our study revealed between- and within-age variabilities in PathoClock and PhysioClock, as well as between-strain variabilities.

Considering organ-specific aging in mouse strains and heterogeneity in organ aging in humans, it is of paramount importance to disentangle individual-specific and organ-specific aging and how each disease state and adaptation state incorporates into the whole-body system as a function of BODN. The PathoClock and PhysioClock can be employed as translatable tools, recapitulating human Body Clock. These clocks can be used across various species and in both males and females to determine common and distinguished pathologies and physiological assessments applied to age-related healthspan. Quantifying individual Clock levels can be used to more precisely understand mechanisms of aging [41–43] and assess the rate of aging using cross-species translational tools to disentangle age-related similarities and differences and assess organ-, strain- and sex-specific effects of aging intervention studies. Using Bayesian inference allows us to predict such Clocks in established as well as new models and updates can be made when new information at physiological or pathological data become available.

## Acknowledgment

Supported by NIH grants NIA-B K01AG059898 (PI, Shabnam Salimi), R24 AG047115, R56 AG058543, and R01 AG057381 (PI, Warren Ladiges), and P30 AG067000 (PI, Kaberlein).

## Disclosure

No conflicts of interest.

## Notes

### Competing Interest Statement

The authors have declared no competing interest.

